# The transcriptional regulator SpxA1 influences the morphology and virulence of *Listeria monocytogenes*

**DOI:** 10.1101/2022.05.31.494264

**Authors:** Monica R. Cesinger, Oluwasegun I. Daramola, Lucy M. Kwiatkowski, Michelle L. Reniere

## Abstract

*Listeria monocytogenes* is a gram-positive facultative anaerobe and an excellent model pathogen for investigating regulatory changes that occur during infection of a mammalian host. SpxA1 is a widely conserved transcriptional regulator that induces expression of peroxide-detoxifying genes in *L. monocytogenes* and is thus required for aerobic growth. SpxA1 is also required for *L. monocytogenes* virulence, although the SpxA1-dependent genes important in this context remain to be identified. In this work, we sought to investigate the role of SpxA1 in a tissue culture model of infection and made the surprising discovery that Δ*spxA1* cells are dramatically elongated during growth in the host cytosol. Quantitative microscopy revealed Δ*spxA1* also forms elongated filaments extracellularly during early exponential phase in rich medium. Scanning and Transmission Electron Microscopy (STEM) analysis found the likely cause of this morphological phenotype is aberrantly placed division septa localized outside of cell midpoints. Quantitative mass spectrometry of whole cell lysates identified SpxA1-dependent changes in protein abundance, including a significant number of motility and flagellar proteins that were depleted in the Δ*spxA1* mutant. Accordingly, we found that both the filamentation and the lack of motility contributed to decreased phagocytosis of Δ*spxA1* by macrophages. Overall, we identify a novel role for SpxA1 in regulating cell elongation and motility, both of which impact *L. monocytogenes* virulence.

**IMPORTANCE:** *Listeria monocytogenes* is an environmental saprophyte and the causative agent of the serious foodborne disease listeriosis. *L. monocytogenes* uses a highly complex transcriptional network to rapidly respond to the transition from living in the environment to infecting a host. One of the regulators important for *L. monocytogenes* pathogenesis is SpxA1, although the SpxA1-regulated genes necessary for virulence remain undefined. This study set out to examine the role of SpxA1 during infection and discovered that SpxA1 regulates cell shape and motility during both extracellular and intracellular growth. Together, our analyses find that SpxA1 has a multifaceted role in virulence, as SpxA1 regulation of both cell morphology and motility are important for infection.

## INTRODUCTION

*Listeria monocytogenes* is the causative agent of the foodborne disease listeriosis and a well-described model intracellular pathogen (1). *L. monocytogenes* lives as an environmental saprophyte until consumed by a susceptible mammalian host. Once ingested, *L. monocytogenes* is phagocytosed by professional phagocytes or induces its own uptake into non-phagocytic cells (2). Quickly after internalization, *L. monocytogenes* escapes the vacuole and replicates within the host cytosol (3). Here, *L. monocytogenes* produces the surface-associated protein ActA which recruits host actin to mediate intracellular motility and intercellular spread via characteristic actin comet tails (4). The asymmetric distribution of ActA along the bacterial cell surface is tightly coupled to the bacterial cell cycle and critical for motility initiation, comet tail formation, and bacterial movement within the host cytosol (5). While the complex set of biophysical steps necessary for efficient actin-dependent cytosolic motility of *L. monocytogenes* are not fully understood, the rod shape of this bacterium is thought to be essential to this process (6, 7).

Alterations to cell shape can have detrimental effects for many bacterial pathogens, as cell shape dictates motility, surface protein localization, innate immune activation, adherence, and invasion into host cells. For example, the helical shape of the foodborne pathogens *Helicobacter pylori* and *Campylobacter jejuni* is critical for flagellar motility in mucus and host colonization (8–10). Similarly, the curvature of the comma-shaped pathogen *Vibrio cholerae* enhances colonization of the small intestine (11). In *L. monocytogenes*, mutants that form chains due to septation defects are deficient for invasion and virulence (12, 13). Further, the shape and geometry of *L. monocytogenes* influence actin tail formation and function while the mechanical stress of actin-based motility effects cell elongation and division (5, 13, 14).

Alternate morphologies can also provide survival advantages to bacterial pathogens. For example, many pathogenic organisms form filaments *in vivo* that are resistant to both phagocytosis and antimicrobials (15). Filamentation describes the process of continual longitudinal growth without formation of septa between newly replicated chromosomes. Increasing cell envelope volume absent of separation results in cells much longer than their bacillary counterparts (16). Filamentous morphologies are induced in most rod-shaped bacteria by a variety of environmental stressors, such as metabolic changes, DNA damage (the SOS response), or alterations to the stoichiometry of cell division components (15). In addition to these stressors, *L. monocytogenes* filamentation is induced extracellularly in response to high or low pH, salt, cold, and heat stress (17–19). Despite these extracellular observations, the regulation of *L. monocytogenes* morphology *in vivo* and the relevance to pathogenesis is not well understood.

One *L. monocytogenes* regulator that is important for both extracellular and intracellular growth is the redox-responsive transcriptional regulator SpxA1 (20). SpxA1 belongs to the ArsC-like Spx family of proteins that is highly conserved in low G+C Firmicutes (21). *L. monocytogenes* SpxA1 is essential for aerobic growth in rich broth as well as virulence in a murine model of infection (22, 23). Similarly, *Staphylococcus aureus* Spx is essential for growth in rich media and Spx homologues are required for virulence in *Streptococcus* spp. and *Enterococcus faecalis* (24–28). Our previous work found that *L. monocytogenes* lacking *spxA1* are unable to replicate aerobically due to toxic levels of endogenously produced hydrogen peroxide, resulting from a combination of dysregulated cytochrome oxidase and insufficient catalase production (29). However, the SpxA1-regulated genes necessary for aerobic growth are dispensable for infection and the role of SpxA1 in pathogenesis remains to be defined.

In this work we sought to investigate the role of SpxA1 in *L. monocytogenes* virulence and made the surprising discovery that Δ*spxA1* mutant cells exhibit variable and elongated morphology during growth in the host cytosol. Quantitative microscopy further revealed Δ*spxA1* cells undergo filamentation extracellularly in anaerobic rich medium. A whole cell proteomics approach identified SpxA1-dependent changes in protein abundance, including motility and flagellar proteins that were depleted in the Δ*spxA1* mutant. Finally, we show that filamentation and decreased motility of Δ*spxA1* contribute to the virulence defect of Δ*spxA1* in tissue culture models of infection. Together, these results significantly advance our understanding of the role of SpxA1 in *L. monocytogenes* replication and pathogenesis.

## RESULTS

### *L. monocytogenes ΔspxA1* exhibits an elongated morphology

The *L. monocytogenes* Δ*spxA1* strain is significantly impaired for intracellular replication and intercellular spread, although the SpxA1-regulated genes important for these processes have yet to be identified (23, 29). To further examine the role of SpxA1 in *L. monocytogenes* virulence, we took a microscopy-based approach and infected monolayers of murine fibroblasts grown on glass coverslips. Monolayers were fixed 10 hours post-infection and *L. monocytogenes*, DNA, and host actin were fluorescently labeled. As expected, wild type (wt) *L. monocytogenes* appeared as short rods in the host cytosol. Many bacteria colocalized with increased densities of host actin (red), and some appeared to be forming canonical actin comet tails (Fig 1A). In contrast, intracellular Δ*spxA1* were more heterogeneous in shape, with some appearing as short rods and many appearing as chains of cells or elongated single cells. Furthermore, elongated Δ*spxA1* appeared to colocalize less frequently with highly dense areas of actin. Those cells that did colocalize with host actin displayed disorganized or non-polar actin recruitment and were rarely associated with actin comet tails (Fig 1B). These results led us to question if the morphological defect of Δ*spxA1* was specific to growth in the mammalian cytosol.

**Figure 1.**
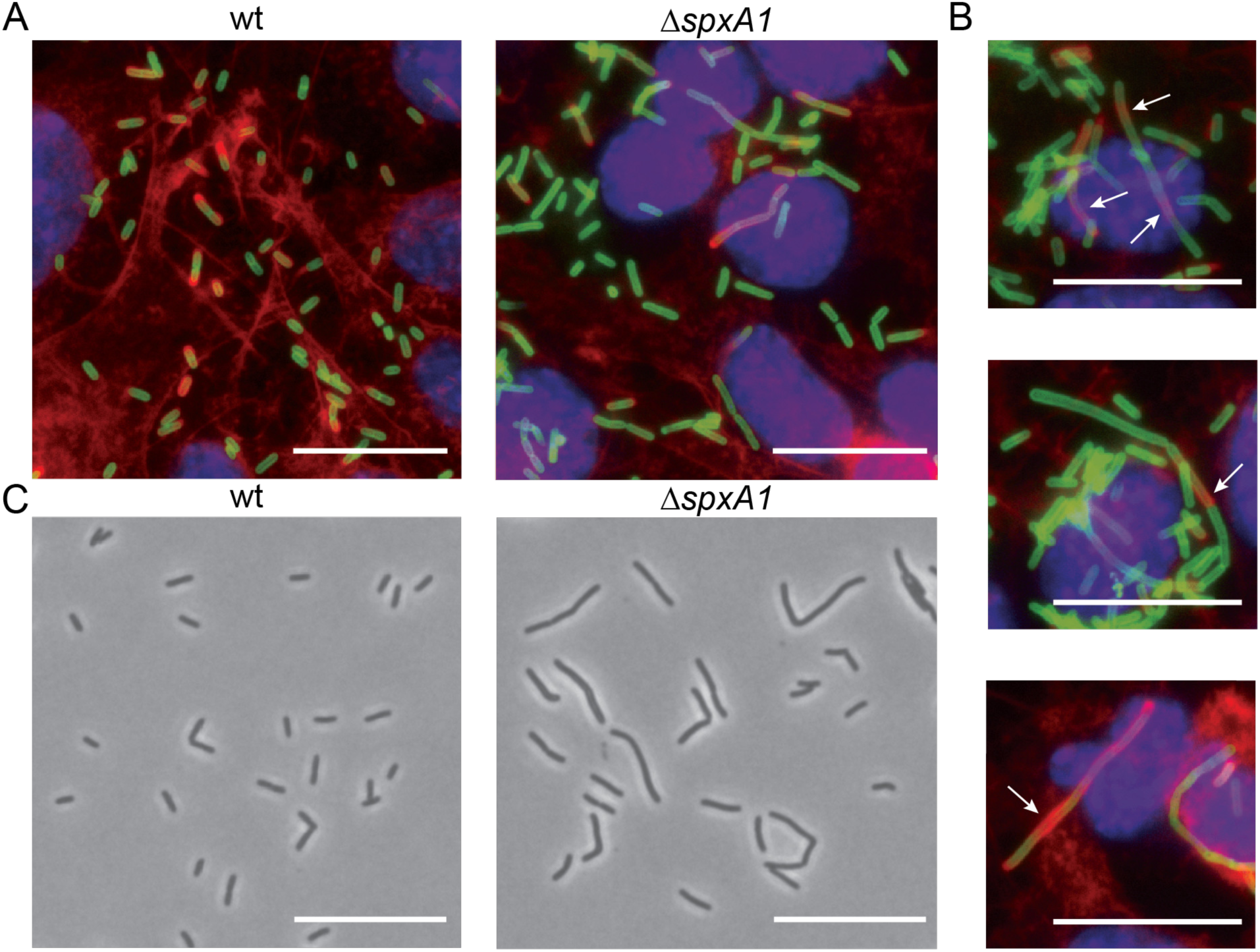
Microscopy of *L. monocytogenes* Δ*spxA1* during intracellular and extracellular growth. (A) L2 murine fibroblasts were infected for 10 hours with wt or Δ*spxA1* and fluorescently labeled to visualize DNA (blue), *L. monocytogenes* (green), and host actin (red). Images were taken using a 100x objective. (B) Additional examples of cells infected with Δ*spxA1.* White arrows indicate elongated bacteria with disorganized, non-polar, or lack of actin recruitment. (C) Phase contrast images of wt and Δ*spxA1* grown to early exponential phase in rich anaerobic broth. Bacteria were imaged with a 40x objective. All scale bars represent 10 µm and all images are representative of three biological replicates and at least ten fields of view.

To determine if the observed Δ*spxA1* morphological heterogeneity was specific to intracellular growth, bacteria were grown to early exponential phase in rich media (brain heart infusion, BHI) in an anaerobic chamber and spotted onto PBS agar pads for phase contrast microscopy. In these conditions, wt *L. monocytogenes* were almost exclusively rod-shaped and of uniform length, whereas the Δ*spxA1* mutant exhibited varied and elongated morphologies (Fig 1C). Together, these images suggested that SpxA1 regulation of cell shape could play a significant role in *L. monocytogenes* growth in both intracellular and extracellular environments.

### Quantification of Δ*spxA1* elongation

To quantitively evaluate Δ*spxA1* morphology, phase contrast images of wt and Δ*spxA1* grown in BHI were analyzed with the cell segmentation software Celltool (30). Cell perimeter was used as the primary metric to describe cell size and account for the irregular shape and curvature of some Δ*spxA1* cells. Despite demonstrating no significant difference in growth rate compared to wt (Fig S1), Δ*spxA1* cells had an increased mean, median, range, and variance of cell perimeters (Fig 2A). The cell perimeters of wt and Δ*spxA1* differed most at early exponential phase, with the mean of wt cells measuring 8.5 µm compared to 16.9 µm for Δ*spxA1* cells. In addition, the maximum cell perimeter observed for the Δ*spxA1* mutant was 146 µm, nearly triple the size of the largest wt cell. While the average cell perimeter of both wt and Δ*spxA1* decreased over time, Δ*spxA1* cells were not restored to wt size at any point during the eight hour time course. In order to accurately assess the biological relevance of the observed elongation, we calculated the percentage of elongated bacteria. Elongated cells were defined as having a perimeter greater than one standard deviation (SD) above the mean of wt cells at each time point. Approximately 10% of wt cells were elongated at each time point examined (Fig 2B). Throughout growth, Δ*spxA1* cultures contained a much higher percentage of elongated cells, with elongated cells comprising 54% of the population at 2 hours. This phenotype was rescued via genetic complementation by providing a copy of *spxA1* with its native promoter at an ectopic site on the chromosome using the integrative plasmid *pPL2* (Δ*spxA1* pPL2.*spxA1*) (31). These data demonstrated that *L. monocytogenes* lacking *spxA1* form significantly longer cells, particularly at early exponential phase.

**Figure 2.**
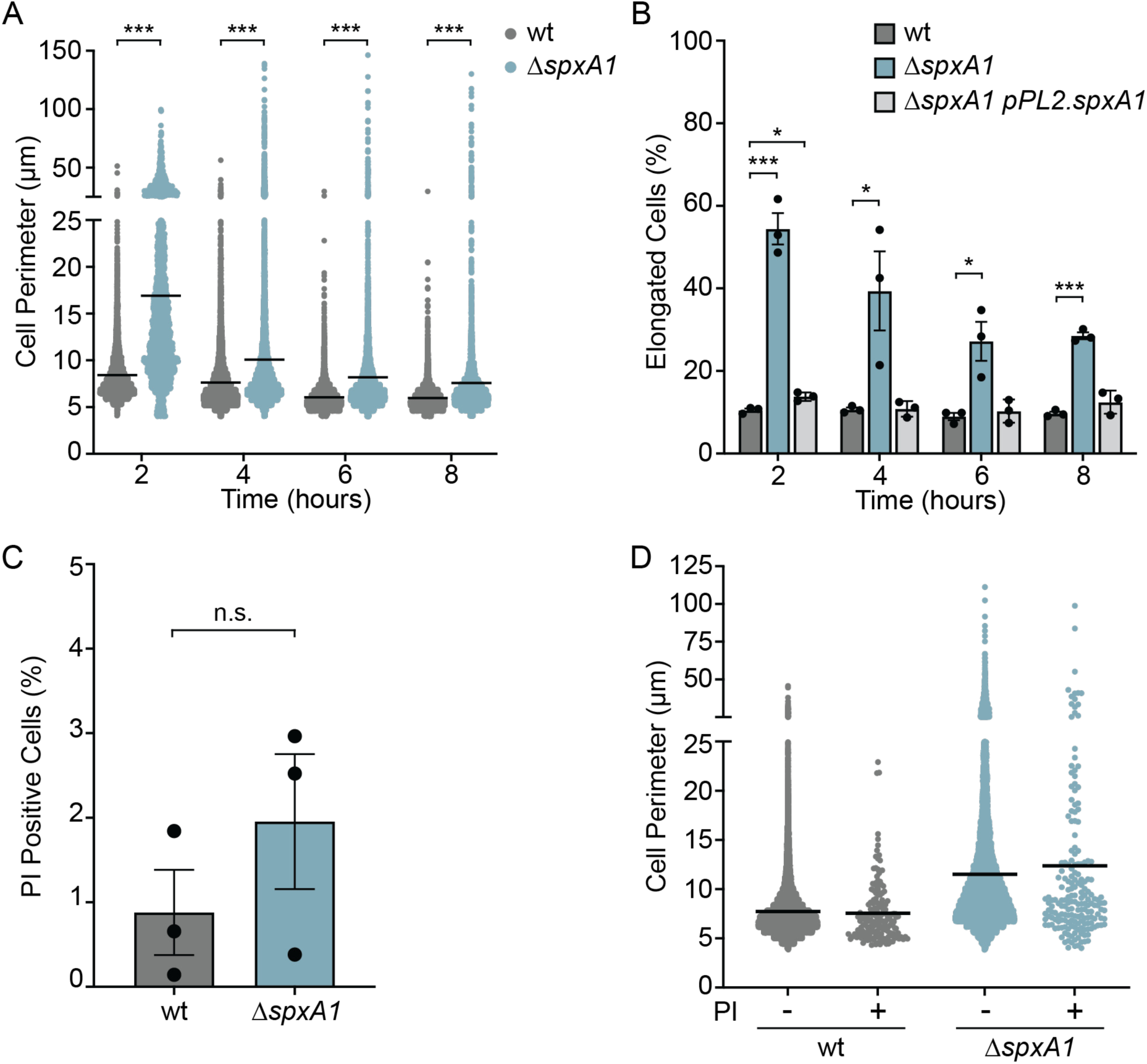
Quantification of *L. monocytogenes ΔspxA1* elongation and cell viability during broth growth. (A) Bacteria were grown anaerobically in rich broth and cell perimeters were measured from phase contrast images using Celltool. The lines indicate the means. Distributions were compared using Kolmogorov-Smirnov tests. *** *p* < 0.001. (B) The percentage of elongated cells was measured for at least ten fields of view or 1,000 cells. Elongated cells were defined as perimeter lengths greater than one standard deviation above the wt mean at each time point. Each symbol represents a biological replicate, the bars indicate the mean, and the error bars denote the standard error of the mean (SEM). *p* values were calculated using an unpaired Student’s *t* test compared to wt. * *p* < 0.05; *** *p* < 0.001. (C) The percentage of propidium iodide (PI) positive cells was calculated from at least 1,000 cells or ten fields of view. The data are not significantly different, as determined by an unpaired Student’s *t* test (n.s.). (D) The cell perimeters of bacteria from PI-negative (-) and PI-positive (+) populations were measured. Graphs represent cell perimeters measured from three biological replicates where at least 1,000 cells or ten fields of view were quantified per replicate. The lines indicate the means.

Due to the observed morphological irregularities, we hypothesized that highly elongated Δ*spxA1* cells may have irregular or damaged membranes and therefore decreased viability. To determine the viability and membrane integrity of the Δ*spxA1* mutant, early log phase cultures were stained with propidium iodide (PI), a live-dead stain which cannot penetrate intact lipid bilayers. We observed an approximate 2-fold increase in the percentage of PI-positive Δ*spxA1* cells compared to wt, although this difference was not statistically significant (Fig 2C). Neither strain exhibited significant PI staining, with the percent of PI-positive cells never reaching 2% for wt or 3% for Δ*spxA1*. To evaluate the viability of elongated cells specifically, the cell perimeters of the PI-positive and PI-negative populations were measured for each strain. The distributions of PI-positive and PI-negative Δ*spxA1* cells appeared nearly identical, indicating that the elongated Δ*spxA1* cells were not more likely to be PI-positive (Fig 2D). Taken together, these results demonstrated that Δ*spxA1* cells are morphologically heterogenous and a significant portion of the cells are dramatically elongated. However, this elongation does not correlate to a critical defect in viability or membrane integrity.

### Fluorescence microscopy of membranes

Phase contrast microscopy revealed that the majority of Δ*spxA1* cells were significantly elongated compared to wt, but the quantitative analysis of these images could not distinguish between incompletely separated chains of cells or filamentous single cells. To examine the specific nature of elongated Δ*spxA1* cells, bacteria were labeled with the fluorescent membrane dye TMA-DPH, which intercalates into lipid bilayers containing fatty acyl chains (32). *L. monocytogenes* wt cells appeared as typical rods at various stages of division with sites of septation localized to the cell midpoint (Fig 3A). In contrast, Δ*spxA1* exhibited a range of elongated filamentous morphologies which rarely contained septa at cell midpoints and often exhibited a complete lack of septa. These results suggested that the significant elongation of the Δ*spxA1* mutant was likely the result of filamentation rather than chaining.

**Figure 3.**
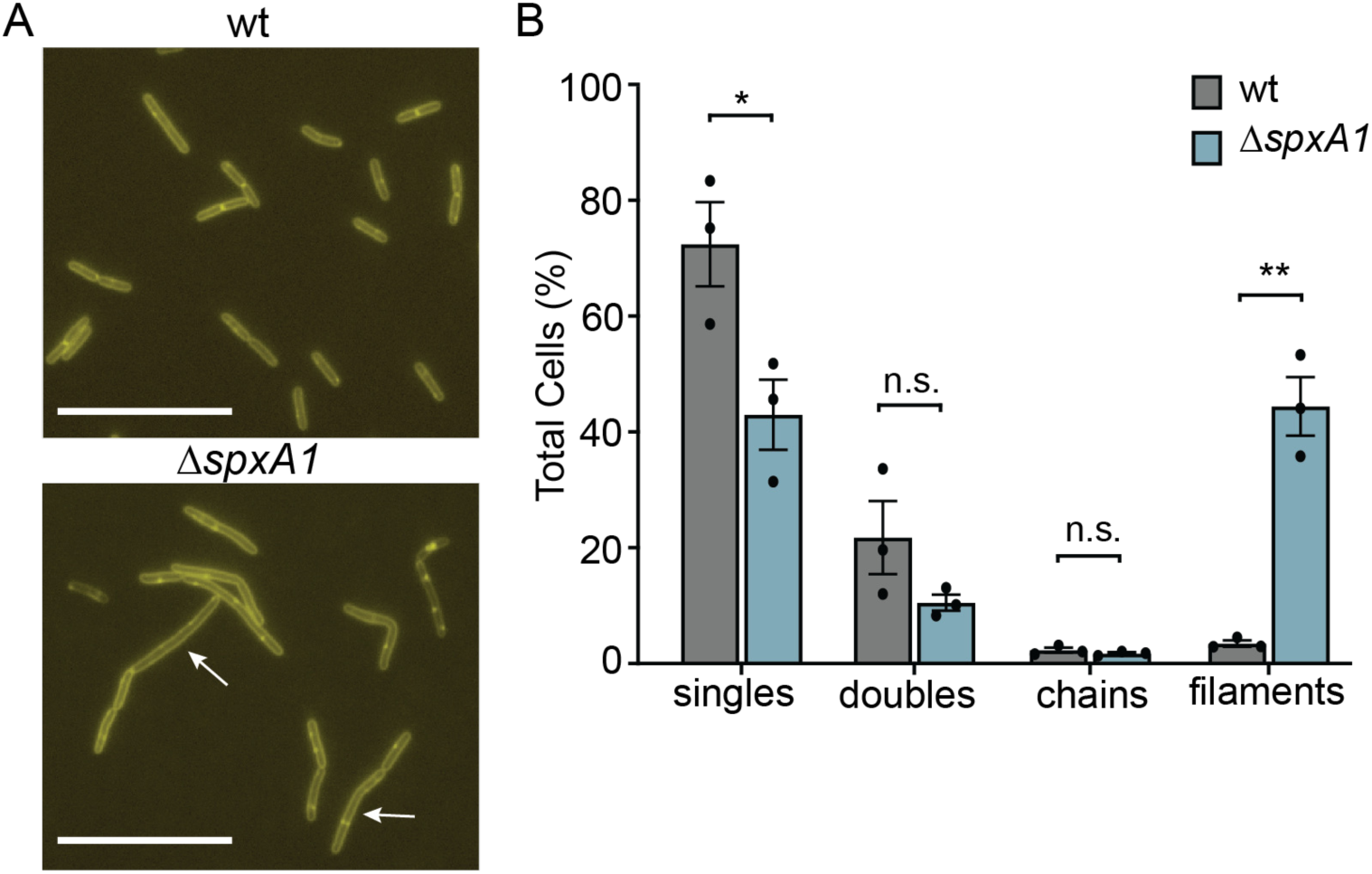
Fluorescent labeling of membranes indicates the prevalence of Δ*spxA1* filamentation. (A) *L. monocytogenes* wt and Δ*spxA1* grown to early exponential phase were labeled with the fluorescent membrane stain TMA-DPH. Arrows indicate elongated cells without regular septa and scale bars represent 10 µm. (B) Quantification of elongated cells observed in *A*. Data displayed are the means and SEM of three biological replicates of 1,000 cells each. *P* values were calculated using an unpaired Student’s *t* test compared to wt. * *p* < 0.05; ** *p* < 0.01; n.s. not significant.

To quantify the frequency of Δ*spxA1* filamentation, we visually evaluated the TMA-DPH images and categorized over 1,000 cells per replicate for each strain. Single cells were defined as cells lacking fully formed septa that were approximately the length of the wt mean (2-5 µm). Doublets and chains were defined as two or more linked bacteria, respectively, composed of 2-5 µm cells attached by evenly placed septa. Filaments were defined as cells longer than 2-5 µm that either lack septa completely, have irregular numbers of septa relative to the length of the cell, or contain septa placed outside the midpoint of the cell. As expected, approximately 94% of wt bacteria were either single cells or doublets, with a small minority of chains and filaments (Fig 3B). In contrast, over 44% of Δ*spxA1* cells were elongated filaments with aberrant septum localization. These results confirmed that the majority of elongated Δ*spxA1* are filamentous single cells.

### Scanning Transmission Electron Microscopy

After determining that the majority of Δ*spxA1* cells were filaments, we next sought to investigate the observed differences in membrane architecture at higher resolution. To that end, we performed Scanning Transmission Electron Microscopy (STEM) on wt, Δ*spxA1*, and the complemented Δ*spxA1* strain. Early exponential phase cultures were negatively stained with uranyl acetate, which incorporates into organic materials with a high affinity for phosphorylated molecules, enabling resolution of dense and highly phosphorylated components, such as membranes, nucleic acids, and some proteins (33). In many respects, the Δ*spxA1* cell envelope appeared indistinguishable from that of wt and the complemented strain (Fig 4). For example, all three strains had a similarly thick outer layer of peptidoglycan covering a thin layer of highly contrasting phospholipid membrane, with no obvious differences in phospholipid or peptidoglycan abundance (Fig 4A). Additionally, the division septa appeared to have similar architecture and composition in all three strains. However, several differences in the spatial organization of the Δ*spxA1* envelope were observed. Consistent with our phase contrast and fluorescence microscopy images, Δ*spxA1* cells appeared elongated compared to wt (Fig 4B-D). Multiple septa were observed at various stages of development within Δ*spxA1* filaments, and these were unevenly distributed along the cell (Fig 4C). We observed Δ*spxA1* cells in the process of initiating septum formation in aberrant locations outside the cell midpoint, sometimes irregularly spaced and atypically close together (Fig 4B and C). Formation of septa was not only initiated, but successfully completed outside of cell midpoints (Fig 4D). Finally, Δ*spxA1* cells could initiate full separation of segmented cells at sites of mis-localized septation (Fig 4D). Taken together, these images suggested that Δ*spxA1* filamentation is due to both improper localization and frequency of septum formation.

**Figure 4.**
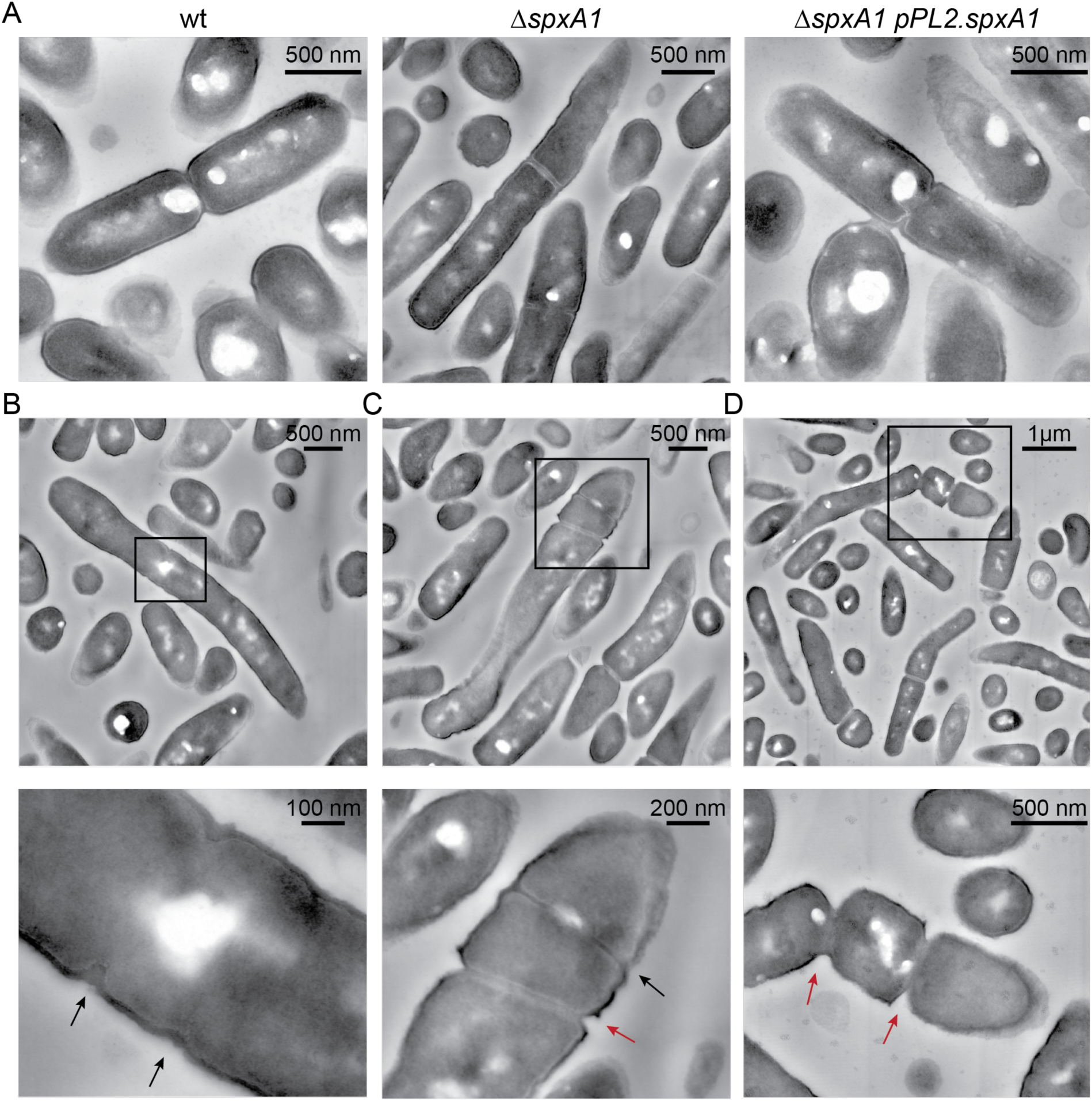
Scanning Transmission Electron Microscopy (STEM) of *L. monocytogenes*. (A) *L. monocytogenes* strains grown to early exponential phase were fixed and imaged by STEM. (B-D) Representative examples of Δ*spxA1* filaments. Boxed portions of each image are shown magnified below. Scale bars are indicated in each panel.

### Whole cell proteomics

To identify SpxA1-dependent factors influencing cell morphology, we performed whole cell proteomics on early exponential phase bacterial cultures. Previous work demonstrated that SpxA1 impacts the transcription of 217 genes, activating 145 and repressing 72, including several proteins involved in protein degradation (29). Thus, a proteomics approach has the advantage of identifying the proteins that are changed directly via SpxA1 transcriptional regulation and indirectly via SpxA1 regulation of protein turnover. Bacteria were grown in triplicate to early exponential phase and proteins were isolated and quantified using unlabeled mass spectrometry coupled to liquid chromatography. We identified 1,692 proteins from 23,219 peptides, accounting for 60.1% of all protein coding regions. Proteins that differed at least 2-fold from wt abundance (*p* > 0.05), or were absent in one strain and abundant (greater than 5 peptides detected) in the other, were considered for further analysis. Using these criteria, we identified 122 proteins that were significantly decreased in abundance in Δ*spxA1* compared to wt (Table S1), and 46 proteins that were significantly increased in abundance in Δ*spxA1* (Table S2).

The largest category of proteins depleted in Δ*spxA1* compared to wt were proteins involved in redox homeostasis and respiration. Specifically, the largest decrease observed was for the protein catalase (50-fold), which detoxifies hydrogen peroxide and is produced during aerobic growth in an SpxA1-dependent manner (29). Further, proteomics detected thiol peroxidase (Tpx) and heme biosynthesis proteins (HemE and HemH) in wt cells at high abundance, but these proteins were below the limit of detection in Δ*spxA1* (Table S1). Other redox-related proteins depleted in Δ*spxA1* compared to wt included: ChdC (heme peroxidase), Lmo0983 (glutathione peroxidase), Lmo1609 (thioredoxin), Rex (redox-responsive regulator), and the SpxA1 paralogue SpxA2 (Lmo2426), which currently has no known function (23). To test if the depletion of these proteins contributed to the morphology of Δ*spxA1* cells, we measured the cell perimeters of deletion mutants grown to early exponential phase in anaerobic BHI. *L. monocytogenes* cells lacking *kat, tpx, hemEH, chdC, lmo0983, lmo1609, rex,* or *spxA2* were not significantly elongated compared to wt cells (Fig S2). These results indicated that the genes critical for Δ*spxA1* aerobic growth and redox homeostasis are not required for proper cell elongation and division.

We next evaluated protein turnover, as proteins in this category were the most dramatically enriched in Δ*spxA1* compared to wt (Table S2). Specifically, the protease adaptor proteins MecA and ClpE were the most increased in Δ*spxA1* (10- and 19-fold respectively). To explore the relevance of these proteins to *L. monocytogenes* morphology, mutants lacking *mecA* or *clpE* were generated via allelic exchange. Quantitative microscopy of *L. monocytogenes ΔmecA* and Δ*clpE* revealed no change in the frequency of cell elongation compared to wt (Fig 5). Similarly, deletion of *mecA* and *clpE* in the Δ*spxA1* background did not rescue the elongation of Δ*spxA1*. Interestingly, the Δ*spxA1*Δ*clpE* double mutant was significantly more elongated than the Δ*spxA1* parental strain. To extend our analysis of protein turnover, mutants lacking *yjbH* and *clpX* were generated in both wt and Δ*spxA1* backgrounds. SpxA1 abundance is regulated by YjbH- and ClpX-dependent degradation such that mutants deficient for *yjbH* or *clpX* have increased SpxA1 (20, 34). *L. monocytogenes* Δ*yjbH* and *clpX::Tn* mutants formed cells significantly smaller than wt (Fig 5). Furthermore, double mutants that lack *spxA1* and *yjbH* or *clpX* did not differ in elongation compared to the Δ*spxA1* parental strain. Together, these data suggested that cell size maintenance requires precise regulation of SpxA1, as mutants with increased SpxA1 abundance form cells smaller than wt and Δ*spxA1* cells are significantly elongated.

**Figure 5.**
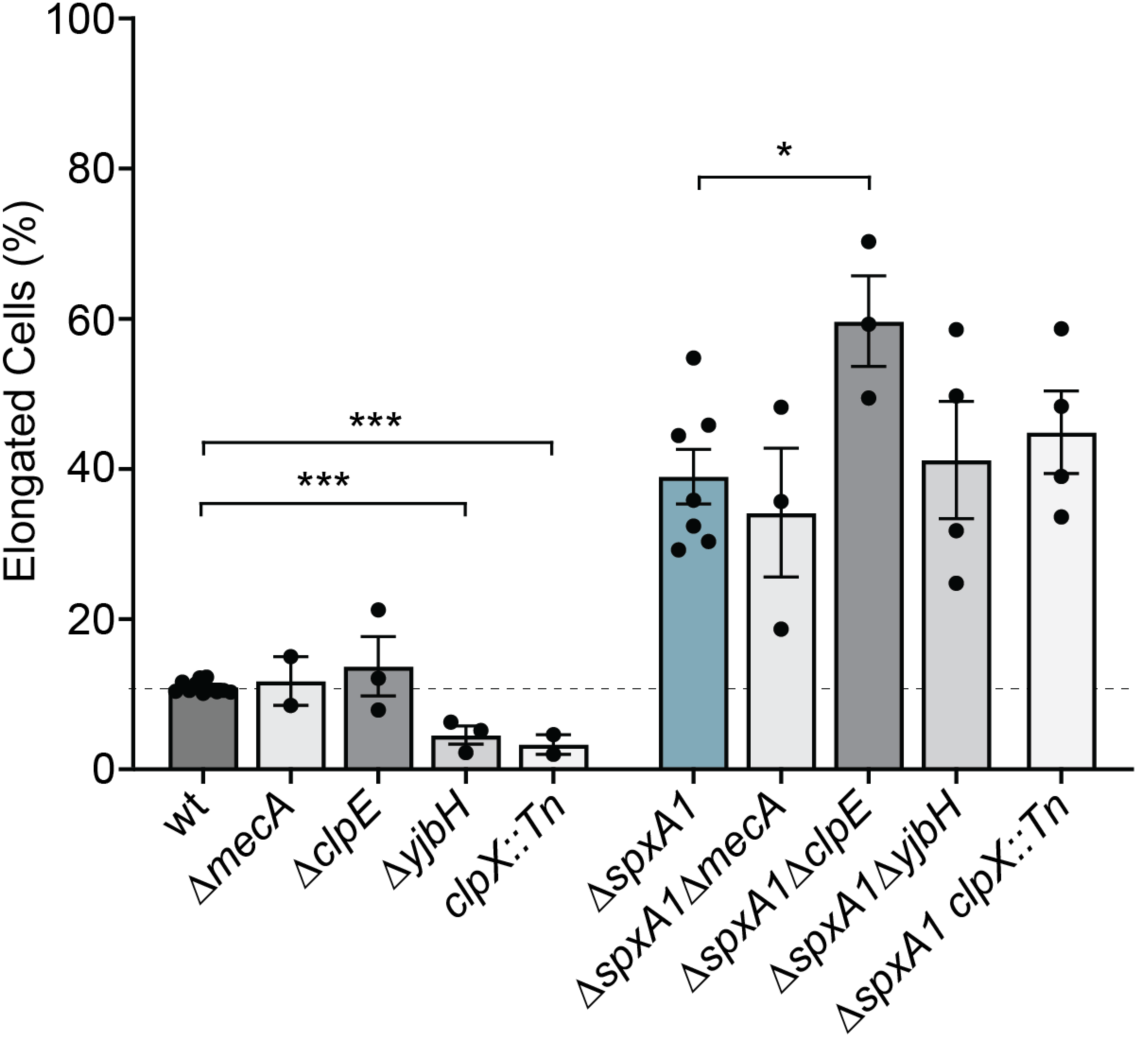
Proteolysis influences morphology of *L. monocytogenes*. The percentage of elongated cells was measured for at least ten fields of view or 1,000 cells. Elongated cells were defined as perimeter lengths greater than one standard deviation above the wt mean. Each symbol represents a biological replicate, the bars indicate the mean, and the error bars denote the SEM. *p* values were calculated using an unpaired Student’s *t* test comparing to the parental strain, as indicated. * *p* < 0.05; *** *p* < 0.001.

We next examined whether *spxA1* elongation was mediated by well-described inducers of filamentation in other Firmicutes. Filamentation can arise due to changes in the abundance of proteins involved in cell envelope biosynthesis and modification, induction of the DNA damage response (the SOS response), or alterations to the stoichiometry of cell division proteins (15). Interestingly, none of the canonical factors critical for these processes were significantly changed in Δ*spxA1* compared to wt. Specifically, proteins involved in the synthesis and modification of wall teichoic acid (DltACD, TagABDH), lipoteichoic acid (LtaPS, LafAB, GtlAB) and peptidoglycan (MurBCDEFGIZ, PBPs, Ami, PgdA) were equally abundant in wt and Δ*spxA1* (Table S3). Furthermore, immunoblot detection of lipoteichoic acid revealed no changes in the size or composition of lipoteichoic acid between wt and Δ*spxA1* (data not shown). The SOS response induces filamentation in response to DNA damage through RecA-mediated transcriptional changes. RecA binds to single-stranded DNA and induces the auto-cleavage of the major SOS regulator LexA, leading to inhibition of septation (35). Whole cell proteomics identified no difference in the abundance of RecA or LexA in Δ*spxA1* compared to wt, and several other canonical SOS proteins were also unchanged (RecF, RecQ, RuvAB, and UvrABC). Lastly, critical cell shape determinants (MreBC, RodA1, RodZ), proteins necessary for the formation of the divisome (FtsAEHKYZ, SepF, ZapA,), and proteins required for septum localization to the cell midpoint (DivIB, EngB, EzrA, MinCDJ, Noc) were unchanged in the Δ*spxA1* mutant compared to wt. Together, we did not find evidence of cell envelope alterations, induction of the SOS response, or aberrant production of divisome proteins in Δ*spxA1* filamentous cells.

Finally, we examined whether Δ*spxA1* filamentation was the result of a deficiency in secreted factors such as autolysins, small peptides, or signaling molecules that would not be readily measured by whole cell proteomics. If this was the case, we hypothesized that co-culture with wt would rescue Δ*spxA1* filamentation *in trans*. Indeed, *L. monocytogenes* mutants lacking the secreted peptidoglycan hydrolases NamA or p60 form chains that are unable to divide unless exposed to wt culture supernatants (12, 13). However, anaerobic co-culture of wt constitutively expressing GFP with Δ*spxA1* expressing mCherry did not rescue the elongated phenotype of Δ*spxA1* (Fig S3). Overall, our proteomic and microscopic analyses determined that SpxA1-dependent factor(s) regulate cell size, as mutant cells with increased SpxA1 are significantly smaller than wt and *L. monocytogenes* lacking *spxA1* are significantly elongated. Moreover, we determined that SpxA1-dependent filamentation is not due to SOS response induction, altered synthesis or modification of teichoic acids or peptidoglycan, changes in cell division protein abundance, or defects in secreted factors such as autolysins or small molecules.

### The roles of motility and morphology during infection

Upon examining our whole cell proteomics for protein changes that could impact virulence, we discovered that proteins associated with motility and chemotaxis were significantly decreased in abundance in Δ*spxA1* compared to wt (Table 1). Specifically, Δ*spxA1* was significantly deficient for 16 motility and chemotaxis proteins, including a 7-fold decrease in the flagellin monomer FlaA. We hypothesized that decreased motility contributes to the virulence defect of Δ*spxA1* in tissue culture infection models, as previous research has demonstrated that non-motile mutants are unable to swim towards host cell monolayers and thus cannot efficiently invade (36, 37). To address this hypothesis, gentamicin protection assays were performed on infected immortalized murine bone marrow-derived macrophages (iBMMs) and uptake of Δ*spxA1* was compared to that of wt or a Δ*flaA* mutant that lacks flagella (37). After a 30 minute infection with *L. monocytogenes*, cells were washed and incubated with media containing gentamicin for 30 minutes to kill extracellular bacteria. Host cells were then lysed, plated to enumerate CFU, and intracellular bacteria were calculated as a percentage of the total inoculum. The Δ*flaA* and Δ*spxA1* mutants exhibited similar ∼10-fold reductions in intracellular bacteria compared to wt *L. monocytogenes* (Fig 6A). To test if the Δ*spxA1* defect was due to impaired motility, centrifugation was applied immediately after infection to force the bacteria to contact the host cell monolayer. Following centrifugation, the Δ*flaA* and Δ*spxA1* mutants were taken up by iBMMs with the same efficiency as wt *L. monocytogenes* (Fig 6B). These results suggested that the observed defect in phagocytosis of Δ*spxA1* can be attributed to decreased motility that limits interactions with the host cells.

**Figure 6.**
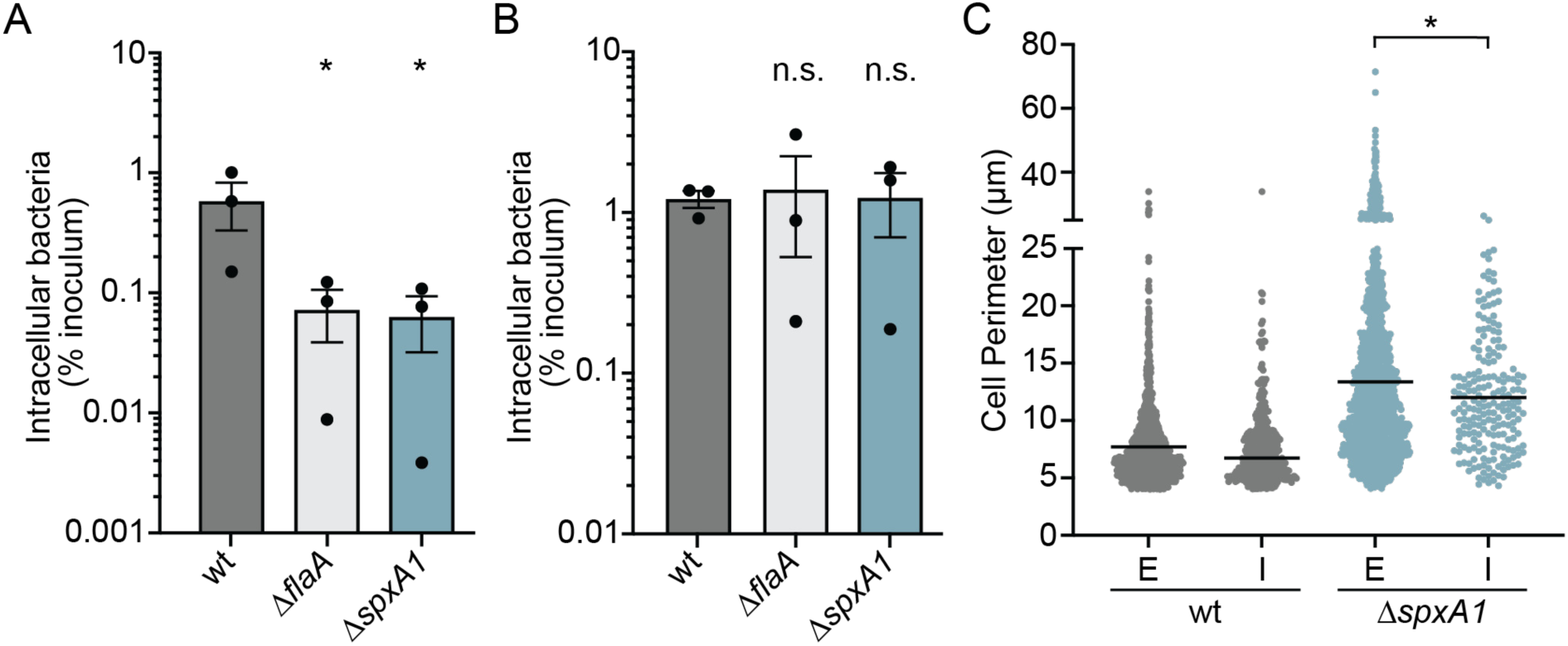
The roles of bacterial motility and morphology in macrophage phagocytosis of Δ*spxA1*. (A) Gentamicin protection assay measuring intracellular bacteria 1 hour post-infection of immortalized bone marrow-derived macrophages (iBMMs). (B) Gentamicin protection assay after bacteria were centrifuged onto the host cells immediately after infection. The mean and SEM of three biological replicates are shown for *A* and *B*. *p* values were calculated using an unpaired Student’s *t* test comparing to wt. * *p* < 0.05; n.s. not significant. (C) iBMMs were infected with centrifugation for 15 minutes and were stained to enable differentiation of extracellular (E) and intracellular (I) bacteria. The perimeters of cells in each compartment were measured by Celltool. Cell perimeters were measured from two biological replicates where at least 1,000 bacteria or ten fields of view were quantified. The lines represent the means. Distributions were compared using Kolmogorov-Smirnov tests. * *p* < 0.05.

**Table 1.**
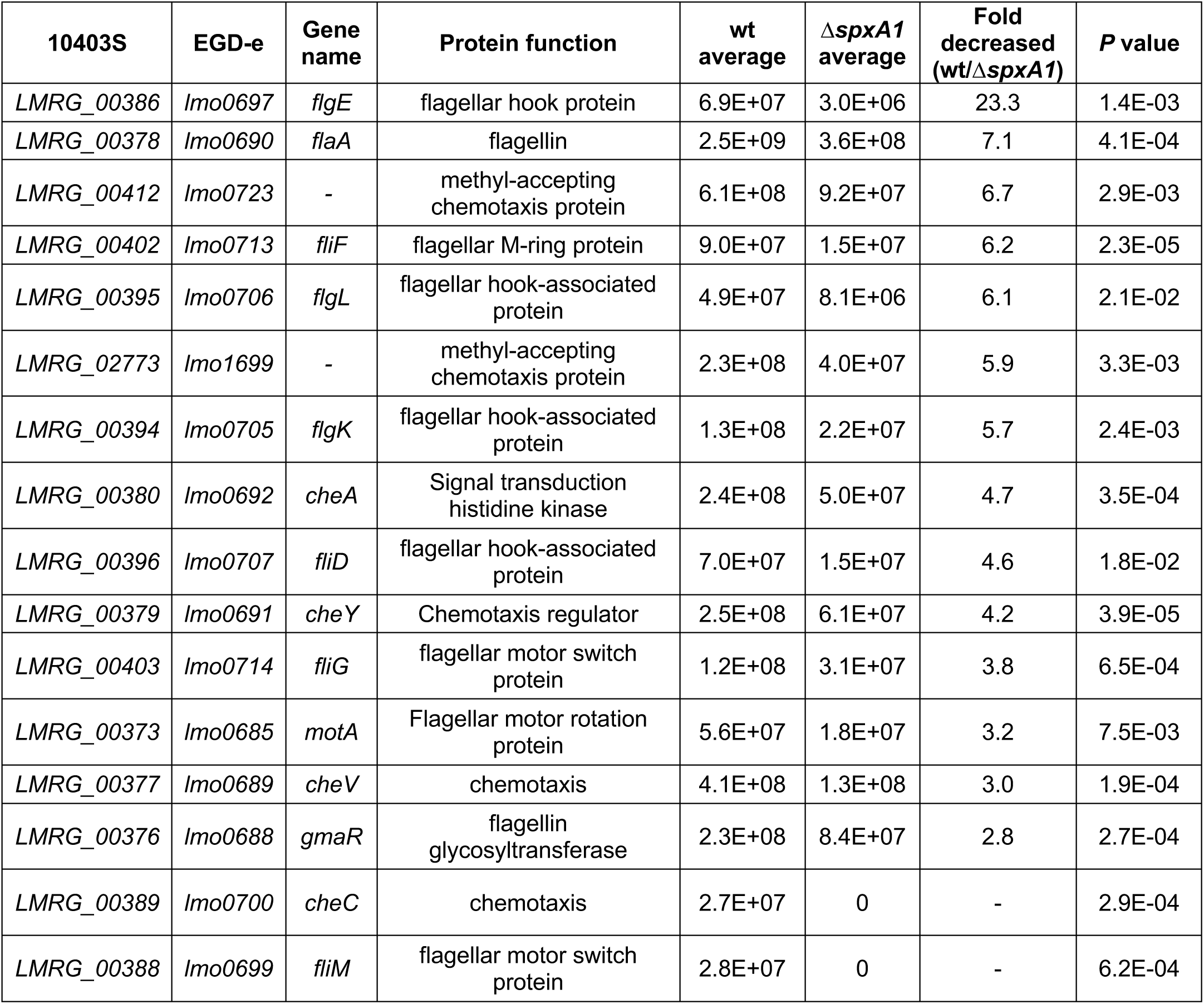
Motility proteins decreased in Δ*spxA1*

| 10403S | EGD-e | Gene name | Protein function | wt average | $\Delta spxA1$ average | Fold decreased (wt/$\Delta spxA1$) | P value |
| --- | --- | --- | --- | --- | --- | --- | --- |
| LMRG_00386 | Imo0697 | *flgE* | flagellar hook protein | 6.9E+07 | 3.0E+06 | 23.3 | 1.4E-03 |
| LMRG_00378 | Imo0690 | *flaA* | flagellin | 2.5E+09 | 3.6E+08 | 7.1 | 4.1E-04 |
| LMRG_00412 | Imo0723 | - | methyl-accepting chemotaxis protein | 6.1E+08 | 9.2E+07 | 6.7 | 2.9E-03 |
| LMRG_00402 | Imo0713 | *fliF* | flagellar M-ring protein | 9.0E+07 | 1.5E+07 | 6.2 | 2.3E-05 |
| LMRG_00395 | Imo0706 | *flgL* | flagellar hook-associated protein | 4.9E+07 | 8.1E+06 | 6.1 | 2.1E-02 |
| LMRG_02773 | Imo1699 | - | methyl-accepting chemotaxis protein | 2.3E+08 | 4.0E+07 | 5.9 | 3.3E-03 |
| LMRG_00394 | Imo0705 | *flgK* | flagellar hook-associated protein | 1.3E+08 | 2.2E+07 | 5.7 | 2.4E-03 |
| LMRG_00380 | Imo0692 | *cheA* | Signal transduction histidine kinase | 2.4E+08 | 5.0E+07 | 4.7 | 3.5E-04 |
| LMRG_00396 | Imo0707 | *fliD* | flagellar hook-associated protein | 7.0E+07 | 1.5E+07 | 4.6 | 1.8E-02 |
| LMRG_00379 | Imo0691 | *cheY* | Chemotaxis regulator | 2.5E+08 | 6.1E+07 | 4.2 | 3.9E-05 |
| LMRG_00403 | Imo0714 | *fliG* | flagellar motor switch protein | 1.2E+08 | 3.1E+07 | 3.8 | 6.5E-04 |
| LMRG_00373 | Imo0685 | *motA* | Flagellar motor rotation protein | 5.6E+07 | 1.8E+07 | 3.2 | 7.5E-03 |
| LMRG_00377 | Imo0689 | *cheV* | chemotaxis | 4.1E+08 | 1.3E+08 | 3.0 | 1.9E-04 |
| LMRG_00376 | Imo0688 | *gmaR* | flagellin glycosyltransferase | 2.3E+08 | 8.4E+07 | 2.8 | 2.7E-04 |
| LMRG_00389 | Imo0700 | *cheC* | chemotaxis | 2.7E+07 | 0 | - | 2.9E-04 |
| LMRG_00388 | Imo0699 | *fliM* | flagellar motor switch protein | 2.8E+07 | 0 | - | 6.2E-04 |

While motility is clearly important for Δ*spxA1* cells to access the host cell monolayer, it is also well-known that filamentous bacteria and fungi are more resistant to phagocytosis than bacilli (15). To test if filamentation contributes to decreased host cell uptake of Δ*spxA1*, we performed quantitative immunofluorescence microscopy of infected iBMMs. *L. monocytogenes* wt or Δ*spxA1* constitutively expressing mCherry were added to cells with centrifugation and allowed to infect for 15 minutes. Cells were then washed, fixed, immunostained with a *L. monocytogenes*-specific antibody, and labeled with a green fluorophore. As the iBMMs were not permeabilized prior to immunostaining, extracellular bacteria could be identified by the colocalization of red and green fluorescence, while intracellular bacteria could be differentiated by the absence of green fluorescence. Celltool was then used to measure the cell perimeters of intracellular and extracellular bacterial populations. We observed a significant decrease in the mean perimeter of intracellular Δ*spxA1* cells compared to extracellular cells and all filamentous Δ*spxA1* cells with a perimeter of 26 µm or larger remained extracellular (Fig 6C). Together, these results suggested that macrophage uptake of Δ*spxA1* is impaired due to both decreased motility and impaired phagocytosis of Δ*spxA1* filaments.

## DISCUSSION

This study set out to define the role of the transcriptional regulator SpxA1 in *L. monocytogenes* pathogenesis. Surprisingly, we observed that Δ*spxA1* exhibits striking morphological heterogeneity during intracellular and extracellular growth, forming single rod-shaped bacteria, chains of cells, and filamentous single cells. Quantitative microscopic analysis of this morphology phenotype *in vitro* revealed that *L. monocytogenes* lacking *spxA1* primarily form filamentous cells with an average cell perimeter twice that of wt. These filamentous cells did not have compromised cell membranes and appeared to result from irregular frequency and localization of division septum formation. Using a whole cell proteomics approach, we determined that SpxA1 does not influence the abundance of proteins critical to cell division, the SOS response, or the synthesis or modification of cell envelope components. However, proteins important for motility and chemotaxis were significantly depleted in Δ*spxA1* compared to wt *L. monocytogenes* and bacterial motility was necessary for infection of macrophages in tissue culture. Furthermore, highly filamentous Δ*spxA1* cells were more resistant to phagocytosis than their bacillary counterparts, supporting a multifaceted role of SpxA1 in virulence regulation. Together, our results provide novel insight into the function of SpxA1 in *L. monocytogenes* and its roles in regulating cell division, motility, and virulence.

To examine the morphological heterogeneity of Δ*spxA1* cells, we took several complimentary microscopy approaches. Phase contrast images revealed that approximately five-fold more Δ*spxA1* cells are elongated in early exponential phase than wt, but these elongated cells did not exhibit a significant loss in viability or membrane integrity. Membrane staining identified that Δ*spxA1* cultures are dominated by filamentous cells, the physiology of which we investigated more closely with STEM. Previous research successfully applied STEM to identify changes in the thickness and composition of peptidoglycan in the *L. monocytogenes* envelope (38). However, our STEM images suggested that the composition of the Δ*spxA1* cell envelope did not differ significantly from wt. Instead, we observed notable irregularities in division septum formation in Δ*spxA1* filamentous cells. Overall, these microscopic analyses suggested that the altered frequency and localization of division septa are likely responsible for Δ*spxA1* filamentation.

To elucidate the mechanism of Δ*spxA1* filamentation, quantitative label-free mass spectrometry of whole cell lysates was used to identify SpxA1-dependent changes in protein abundance. Results from this global proteomic analysis disproved many of our initial hypotheses about the potential mechanism of Δ*spxA1* filamentation. The largest category of proteins influenced by SpxA1 were those involved in redox homeostasis and respiration. However, the SpxA1-regulated redox genes essential for aerobic growth were dispensable for SpxA1 regulation of morphology. Our whole cell proteomics also enforced observations from previous transcriptional analysis that SpxA1 plays a distinct role in regulating components of protein turnover. While the proteins most increased in Δ*spxA1* compared to WT were MecA and ClpE, both related to protein turnover, deleting these proteins did not rescue Δ*spxA1* filamentation. Indeed, the increased filamentation of Δ*spxA1*Δ*clpE* compared to Δ*spxA1* suggests that ClpE over-production in Δ*spxA1* may be a direct response to the disrupted stoichiometries of SpxA1-regulated factors. Our data therefore support a novel model for SpxA1 influence over morphology which is independent of the other previously described roles for this protein.

Whole cell proteomics results excluded the influence of several canonical filamentation pathways on Δ*spxA1* morphology, including the SOS response, alterations to cell division machinery, and cell envelope modifications. Filamentation of bacteria is often associated with induction of the SOS response to DNA damage (15), but key SOS response proteins were unchanged in Δ*spxA1*. The production and modification of cell envelope components can also influence both morphology and pathogenesis in *L. monocytogenes* (39–41). However, proteins required for biosynthesis and modification of teichoic acids and peptidoglycan were not altered in the Δ*spxA1* mutant. Given the septation defects of the Δ*spxA1* mutant, it was most surprising that proteins important for cell division were unchanged in Δ*spxA1* compared to wt. The *Bacillus subtilis* homologue Spx directly regulates the divisome component ZapA, which interacts with the cell division Z-ring protein FtsZ to promote septum formation (42). In *L. monocytogenes*, neither FtsZ nor ZapA were changed in the Δ*spxA1* mutant compared to wt. Overall, we found that Δ*spxA1* filamentation was occurring independently of redox response, SOS response, cell division, and cell envelope biosynthesis. Current research is aimed at elucidating the SpxA1-dependent mechanism of filamentation.

Our data suggest that a combination of motility and morphology defects contribute to reduced phagocytosis of Δ*spxA1*. Whole cell proteomics revealed a significant depletion of proteins involved in motility and chemotaxis in Δ*spxA1* compared to wt. We found that flagellar motility is important for *L. monocytogenes* to interact with the host monolayer during tissue culture models of infection, as forcing the bacteria to contact host cells via centrifugation rescued the defect in uptake by macrophages. Immunofluorescence microscopy of the *L. monocytogenes*-host cell surface interaction revealed that the longest filaments were exclusively extracellular. Together, these results demonstrated roles for both motility and bacterial cell size in macrophage phagocytosis.

It is not yet clear which SpxA1-dependent factors are necessary during murine infection, during which Δ*spxA1* is attenuated over 500-fold in the spleen and 5-logs in the liver (23). Flagellar motility is important for adhesion and invasion of host cells *in vitro*, but dispensable for infection in both oral and intravenous models of murine infection (36, 43). While impaired phagocytosis of filamentous Δ*spxA1* cells may contribute to its virulence defect, entry into host cells is not the only barrier to virulence of Δ*spxA1*. Once in the host cytosol, the Δ*spxA1* mutant has a doubling time of 109 minutes as compared to 47 minutes for wt *L. monocytogenes* and is severely defective for intercellular spread (23). The rod shape of *L. monocytogenes* is required in the host cytosol for establishing the asymmetric actin cloud necessary for canonical comet tail formation and motility (6, 7). Thus, the elongated shape and improper actin polarization observed for Δ*spxA1* filaments may limit cytosolic motility and intercellular spread, which would significantly impair host colonization (3, 44, 45). The impact of disorganized septum formation and filamentation on cytosolic growth, intercellular spread, and systemic infection is an area of ongoing investigation.

This study aimed to identify SpxA1-dependent factors important for *L. monocytogenes* pathogenesis. We discovered that Δ*spxA1* cells form elongated filaments that are impaired for motility and resistant to phagocytosis. While the mechanism of filamentation is not yet known, it is intriguing that the SpxA1-regulated genes necessary for aerobic growth *in vitro* are dispensable during infection and are not involved in cell size regulation or motility. SpxA1-dependent regulation thus independently impacts many areas of *L. monocytogenes* physiology. We predict that the complexity of SpxA1 regulation likely results from selective pressure to respond to a variety of signals in diverse environments. Future research will elucidate the cues that influence SpxA1 activity and the genes it regulates during both extracellular growth and pathogenesis.

## MATERIALS AND METHODS

### Bacterial strains and culture conditions

The bacterial strains used are listed in Table S4. All *L. monocytogenes* strains were derived from the 10403S background and cultured in degassed brain heart infusion (BHI) at 37°C in an anaerobic workstation (Don Whitley Scientific), unless otherwise described. All chemicals were purchased from Sigma Aldrich unless otherwise stated. The following concentrations of antibiotics were used: streptomycin, 200 µg/mL; chloramphenicol, 10 µg/mL (*E. coli*), 5 µg/mL (*L. monocytogenes*); tetracycline, 2 µg/mL; and carbenicillin, 100 µg/mL.

Growth curves were carried out in an anaerobic chamber (10% carbon dioxide, 10% hydrogen, balanced with nitrogen) at 37°C in degassed BHI media. First, colonies were inoculated into broth and grown overnight. Fresh BHI was inoculated with overnight cultures to an OD_600_ of 0.02 in 25 mL in 100 mL flasks and grown statically for 8 hours. Every two hours, cultures were serially diluted with sterile degassed PBS and plated on BHI-streptomycin agar to enumerate CFU.

For co-culture experiments, wt constitutively expressing GFP was mixed with Δ*spxA1* constitutively expressing mCherry in 1:1 or 2:1 wt:Δ*spxA1* ratios in fresh degassed BHI. Cultures were grown to early exponential phase, incubated aerobically for 10 min to activate fluorophores, and spotted onto PBS-agar pads for visualization in phase contrast or fluorescence channels using a 40x high resolution objective.

### Strain construction

Generally, plasmids were introduced into chemically competent *E. coli* strains via transformation. Using *E. coli* SM10 (46), plasmids were transferred into *L. monocytogenes* by conjugation. Alternatively, transducing lysates were generated from *L. monocytogenes* containing desired constructs and used to infect recipient strains. Preparation of lysates was accomplished by mixing a donor strain with U153 phage and incubating overnight in soft agar at 30°C, as described previously (47). Phage were subsequently eluted from agar, filter sterilized, and the resulting lysate was mixed with recipient *L. monocytogenes* for 30 min at room temperature. Transductants were then selected on antibiotic-containing agar at 37°C.

In-frame deletions were constructed using PheS* counterselection of the conjugation-proficient suicide vector pLIM1 (gift from Arne Rietsch, Case Western Reserve University). First, upstream and downstream regions of the gene of interest were amplified, fused via SOE PCR, restriction digested, and ligated into pLIM1. The sequences of all pLIM1-derived vectors were confirmed by Sanger DNA sequencing. Deletion vectors were conjugated into recipient *L. monocytogenes* and integrated into the chromosome. After antibiotic selection, integrants were cured of pLIM1 as previously described (48). Allelic exchange was confirmed by PCR, antibiotic sensitivity, and sensitivity to oxygen. The wt and Δ*spxA1* genomes were fully sequenced and it was determined that these strains differed only at the *spxA1* locus.

Genes were either natively expressed or overexpressed from an ectopic locus using the integrative plasmids pPL1 and pPL2 (31). For complementation, an insert containing a gene and corresponding promoter was PCR-amplified, restriction-digested, and ligated into the appropriate vector. Constitutive expression of fluorophores was accomplished by incorporating the desired gene downstream of the pHyper promoter in pPL2 as previously described (47).

### Tissue culture

L2 murine fibroblasts were generated from L929 cells described previously (22). Immortalized murine bone marrow-derived macrophages (iBMMs) were a generous gift from Joshua Woodward (49). Cell lines were grown at 37°C and 5% CO_2_ in high-glucose Dulbecco’s Modified Eagles Medium (Thermo Fisher Scientific) supplemented with 10% heat-inactivated fetal bovine serum, 2mM sodium pyruvate, and 1 mM L-glutamine. During passaging 100 U/mL of Pen-Strep was added to cell culture media. The day before infection, cells were plated in antibiotic-free media.

### Immunofluorescence microscopy

To visualize intracellular bacteria, L2 fibroblasts were seeded at 1.5 x 10^6^ cells per well in 6-well tissue culture-treated plates containing collagen-coated glass coverslips (Thermo Fisher Scientific) and inoculated with *L. monocytogenes* the next day at an MOI of 50. Bacterial cultures were grown overnight in anaerobic BHI at 37°C and washed twice in PBS prior to inoculation. One hour post-infection, monolayers were washed twice with PBS and media containing 30 µg/ mL gentamicin was added to kill extracellular bacteria. 10 hours post infection, coverslips were washed twice with PBS, fixed for 10 minutes in 4% formaldehyde (Pierce), washed in TBS-Tx (0.1% Triton, 50 mM Tris pH 7.6, 150 mM NaCl), and incubated in antibody buffer (TBS-Tx, 1% BSA) at 4°C overnight. Coverslips were then incubated for 30 minutes with rabbit anti-Listeria O antigen antiserum (Difco) at a concentration of 1:100 in antibody buffer and washed in TBS-Tx followed by a secondary incubation with a 1:200 solution of goat anti-rabbit-Alexa488 (Life Tech) in antibody buffer and Alexa555-Phalloidin (Life Tech) at a concentration of 1:1000. After immunolabeling and washing in TBS-Tx a final time, coverslips were attached to glass slides with ProLong Anti-fade Diamond with DAPI, (Life Tech), left to cure at room temperature overnight, and imaged with a 100x objective.

For fluorescence microscopy of extracellular bacteria, overnight cultures of selected strains were diluted in BHI to an OD_600_ of 0.02 and grown for 2 hours to early exponential phase (OD ∼0.2) in an anaerobic chamber at 37°C. Prior to staining, 1 mL of culture was pelleted at 10,000 x *g* for 1 minute in a tabletop centrifuge and resuspended in PBS which halts further bacterial growth and cell death (23). To label cell membranes, cultures were incubated with 20 µg/mL membrane intercalating agent TMA-DPH for 10 minutes in the dark and then washed twice in PBS prior to spotting on PBS-agar pads. To visualize membrane integrity and viability of bacterial cells, cultures were mixed with 5 µg/mL propidium iodide solution, incubated in the dark for 15 minutes, and then spotted immediately onto agar pads.

All imaging was performed using a Keyence BZ-X710 All-in-one Fluorescence Microscope with either a 40x objective or 100x oil immersion objective and corresponding filter cubes (Keyence). Phase contrast images were taken using bright field settings. Propidium iodide and mCherry were detected using BZ-X Filter TexasRed. TMA-DPH was detected using BZ-X Filter DAPI. GFP and Alexa488 were detected using BZ-X Filter GFP.

### Quantitative analysis of bacterial morphology using Celltool

Phase contrast and fluorescence microscopy was performed on *L. monocytogenes* grown anaerobically shaking in BHI and then resuspended in PBS. Culture resuspensions were immediately spotted on PBS-2% agarose pads, sealed under a glass coverslip and imaged. All quantified images were taken using the 40x objective. Fluorescence images were generated in the appropriate channel for each fluorescent label as specified per experiment. At least 10 fields of view were imaged such that at least 1,000 cells could be analyzed per biological replicate per sample. Binary masks were uniformly applied to each field of view using the MATLAB imbinarize function. Quantitative analysis of binarized images was performed using the Celltool software package (30). Image contours were extracted using the extract_contours functions and smoothed via interpolation to polygons with 100 evenly placed vertices. Fluorescence intensities of labeled bacteria were measured using the extract_images function which specifically extracts intensity per area of fluorescence microscopy images based on contours previously generated from phase contrast images. The measure_contours function was used for the final generation of data output. Perimeter length of contours was used as the primary metric for analysis in order to account for curved and irregular morphologies. Unpaired t-tests were used to measure the difference in perimeter length distributions between populations of bacteria. Percent of elongated cells was calculated by measuring the number of cell perimeters in a population which were greater than one SD above the wt mean for each corresponding replicate and time point. SEM and means were displayed for three biological replicates of percent-elongation analysis.

### Scanning Transmission Electron Microscopy

The appropriate strains were grown to early exponential phase anaerobically in 400 mL BHI. Cultures were centrifuged for 10 minutes at 4,500 RPM, and pellets were washed once in 1M PBS. Pellets of approximately 100 µL volume were mixed with an equal volume of fixative buffer consisting of 4% mass spectrometry grade paraformaldehyde and 0.1 M sodium cacodylate. After 30 minutes, cells were pelleted and mixed with equal volumes fresh fixative buffer for 1 hour at room temperature. Samples were then rinsed with 0.1 M sodium cacodylate buffer, post-fixed in osmium tetroxide buffered with 0.1 M sodium cacodylate for 2 hours, and then washed to remove osmium tetroxide. Samples were then progressively dehydrated with increasing concentrations of ethanol (50%, 70%, 90%, 100%), and twice with 100% acetone. Bacteria were then embedded in 50:50 acetone:resin (Epon) and incubated rotating for 4 hours. Incubation was next performed using 20:80 acetone:resin. After drying for 2 hours, samples were placed in a 100% resin gelatin capsule under light vacuum for 2 hours. Finally, samples were placed in fresh resin molds and incubated at 60°C overnight. Resin blocks were sectioned at 80 nm using a Leica Ultracut 6 Microtome and stained using uranyl acetate and Reynolds lead citrate stains. Samples were imaged with an FEI Tecnai G2 F20 Twin transmission electron microscope at 200 kV.

### Whole-cell proteomics

Protein isolation was performed as previously described (34). *L. monocytogenes* were grown anaerobically in BHI for 2 hrs. Bacteria were pelleted, washed once in PBS, and immediately flash-frozen. Bacterial pellets were resuspended in lysis buffer containing 8M urea and lysed by sonification. Proteins in lysates were reduced via incubation with DTT and alkylated via incubation with iodoacetamide. After quantification of protein concentration via BCA, 300 µg of protein per sample were treated with Trypsin Gold (Promega) overnight. The pH was adjusted to pH 2 using trifluoroacetic acid and then proteins were purified using C18-300 peptide purification columns per manufacturer instructions (Nest Group). After elution, peptides were dried and resuspended in 0.1% formic acid. Autosampler vials were loaded with 100 µL sample at a concentration of 0.5 µg/mL. Mass spectrometry was performed on each sample using an Orbitrap Eclipse Mass Spectrometer (Thermo-Scientific) and liquid chromatography was performed using the Easy-nLC 1000 Liquid Chromatograph (Thermo-Scientific). Raw spectral data was processed using the MaxQuant suite of software with default settings and peptide sequences were searched against the *L. monocytogenes* 10403S reference proteome (UniProt).

The relative abundance of proteins was determined using the LFQ intensity metric generated by analysis with the MaxLFQ algorithm, a generic label-free quantification technology available in MaxQuant (50, 51). The LFQ intensity metric was chosen as it does not depend on normalization with housekeeping proteins for accurate abundance prediction sample-to-sample, relying instead on the assumption that most proteins are minimally changed between experimental conditions. Furthermore, LFQ intensities are the result of calculations using the maximum possible information extracted from samples and retain the absolute scale of original peptide intensities, giving a highly accurate proxy for absolute protein abundance (50). Proteins changed in abundance between wt and Δ*spxA1* were considered for further analysis if they met the following criteria: 1. Proteins differed in average abundance between wt and Δ*spxA1* by at least 2-fold. 2. An unpaired t-test comparing the LFQ intensities of three wt and Δ*spxA1* biological replicates yielded a P-value less than 0.05. 3. To increase the stringency of our hit identification and reduce possible false positives, protein abundance calculations were based on the identification of at least 5 peptides per protein in the sample with higher abundance.

### Measuring phagocytosis via gentamicin protection assay and microscopy

Gentamicin protection assays were performed as previously described (48). Immortalized murine bone marrow-derived macrophages (iBMMs) were seeded in 24-well plates at a density of 6 x 10^5^ cells per well. The next day, overnight anaerobic cultures were washed twice with sterile PBS. Macrophages were infected with an MOI of 10 and the inocula were serially diluted and plated on BHI-strep agar to enumerate CFU. To force bacteria to the cell surface, select 6-well plates were then centrifuged at 300 x g for 2 minutes after inoculation. After 30 minutes, monolayers were washed twice in sterile PBS. New media with 30 µg/mL gentamicin was added for 30 min to eliminate any extracellular bacteria and then monolayers were washed twice and lysed in 250 µL cold 0.1% Triton X-100 in PBS. Lysates were serially diluted and plated on BHI-strep agar to enumerate CFU. Percent uptake was calculated by dividing the number of bacteria recovered from each well by the starting inoculum CFU.

To image uptake of bacteria, iBMMs were seeded in 24-well plates containing ethanol-sterilized glass coverslips at a density of 6 x 10^5^ cells per well. Overnight anaerobic cultures were washed twice with sterile PBS. Macrophages were infected with mCherry-expressing bacteria at an MOI of 10 and centrifuged for 2 minutes at 300 rpm to normalize strain contact with the monolayer. Infections were incubated for 15 minutes and then washed twice in sterile PBS. Coverslips were removed and fixed for 10 minutes in 4% formaldehyde (Pierce), washed in PBS, and blocked in PBS with 1% BSA at 4°C overnight. PBS replaced TBST-Tx for all buffers used to prevent the permeabilization and immunolabel infiltration of host cells. Thus, only extracellular bacteria were labeled. Coverslips where then incubated successively with the described immunostaining agents diluted in PBS with 1% BSA, with PBS washes between incubations: 1:20 Fc Block (BD), 1:100 rabbit anti-Listeria O antigen antiserum (Difco), 1:200 goat anti-rabbit-Alexa488 (Life Tech). After washing with PBS a final time, coverslips were attached to glass slides with ProLong Anti-fade Diamond, (Life Tech), left to cure at room temperature overnight, and imaged with a 100x objective. Celltool was used to extract contours from mCherry images (30). Cells were determined to be extracellular based on the intensity of fluorescence extracted from GFP images within the same area as Celltool-derived contours.

## ACKNOWLEDGEMENTS

The authors would like to thank members of the Woodward (UW) and Reniere labs for helpful discussions and critical reading of the manuscript. We also thank the lab of Dr. Joseph Mougous (UW) for access to reagents and equipment and specifically, Dr. Yaxi Wang and Dr. Andi Liu for their assistance with the processing of whole cell proteomics data. STEM imaging was conducted at the Molecular Analysis Facility (MAF) with the help of Dr. Ellen Lavoie. The MAF is supported in part by funds from the Molecular Engineering & Sciences Institute, the Clean Energy Institute, and the National Science Foundation (NNCI-2025489 and NNCI-1542101). Research in the Reniere lab is funded by NIH R01 A132356. M.R.C. was funded by the Howard Hughes Medical Institute through the James H. Gilliam Fellowship for Advanced Study program (GT11030). The funders had no role in study design, data collection and interpretation, or the decision to submit the work for publication.

---

**Figure S1.**
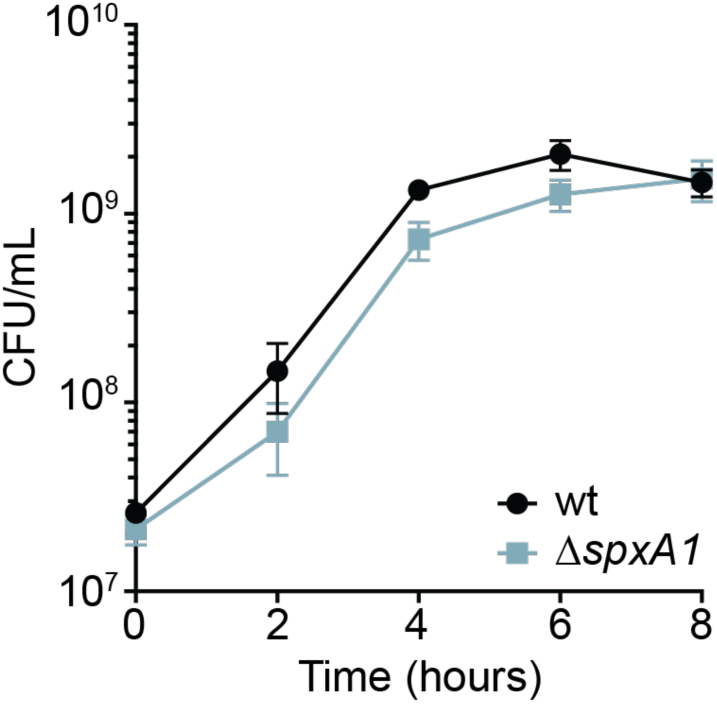
Anaerobic growth kinetics of *L. monocytogenes* wt and Δ*spxA1*. Anerobic cultures were diluted to an OD_600_ of 0.02 in BHI and grown anaerobically. Every two hours, bacterial cultures were serially diluted and plated to enumerate CFU. Data are the means and SEM of three biological replicates. The differences between wt and Δ*spxA1* were not statistically different at any time point, as determined by unpaired t-test.

**Figure S2.**
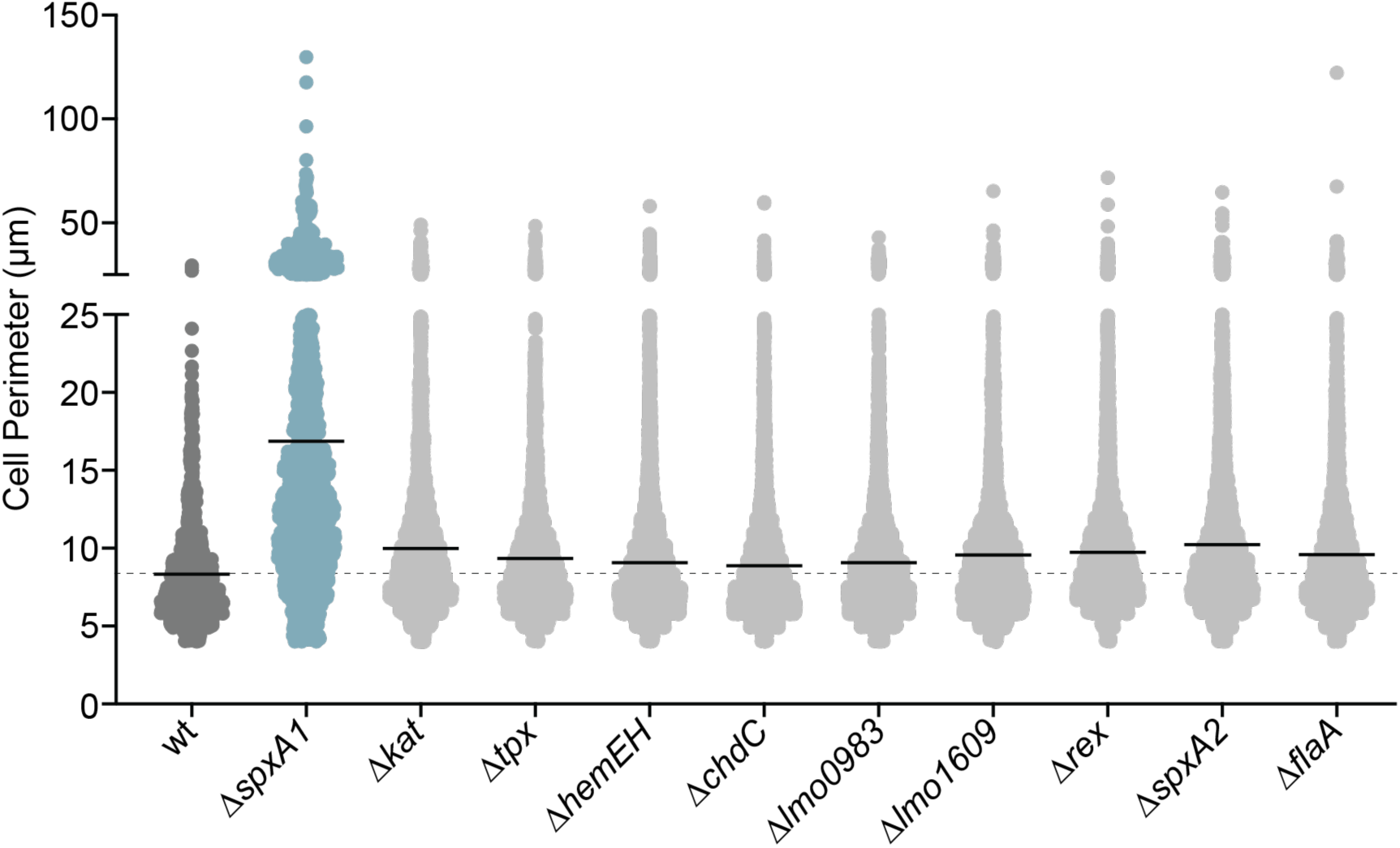
SpxA1-regulated redox homeostasis genes do not influence elongation. Bacteria were grown anaerobically in rich broth and cell perimeters were measured from phase contrast images using Celltool. The lines indicate the means.

**Figure S3.**
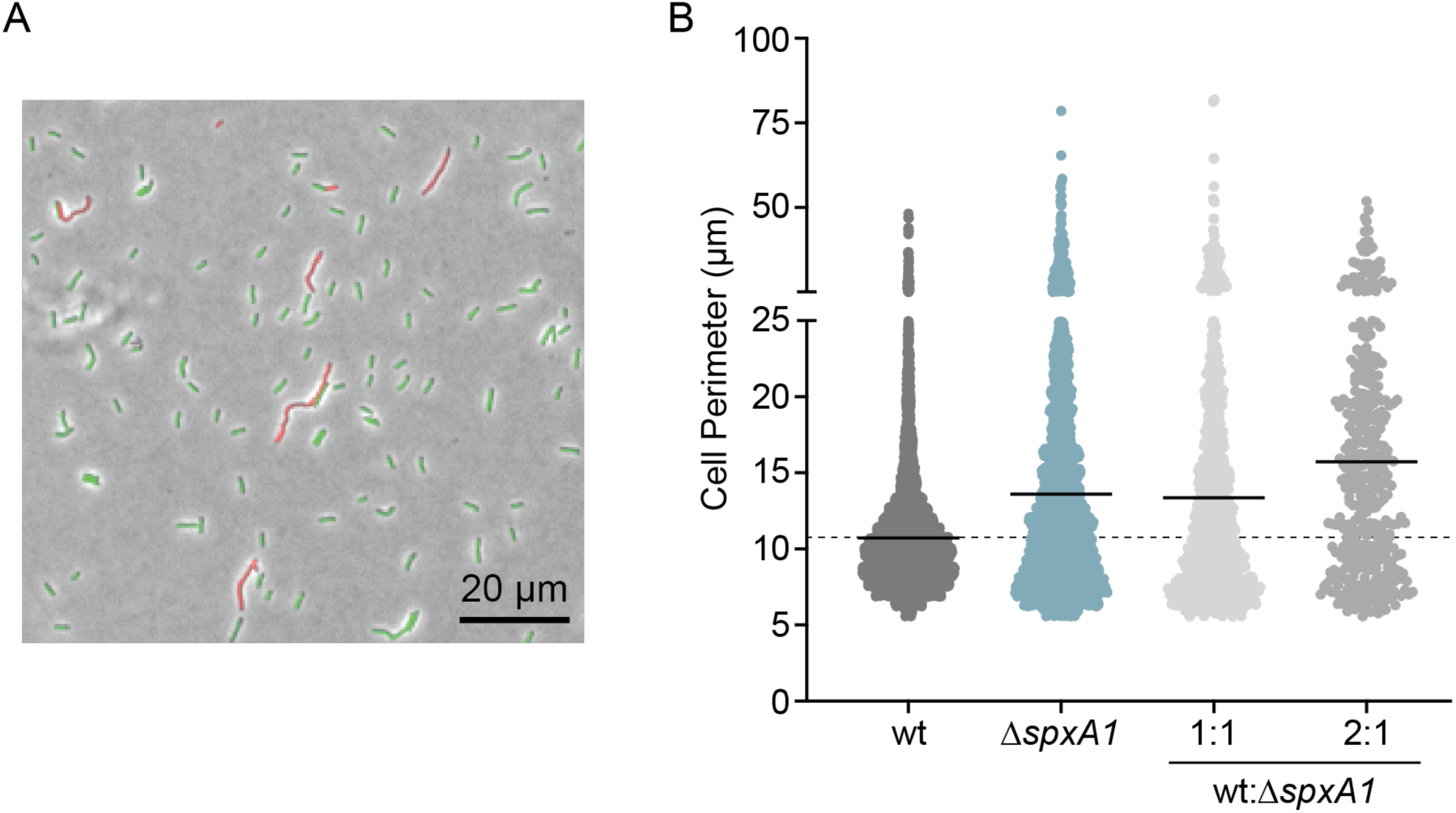
Co-culture with wt does not rescue Δ*spxA1* filamentation. *L. monocytogenes* wt constitutively producing GFP was co-cultured with Δ*spxA1* constitutively producing mCherry for 2 hours. (A) A representative image of a 2:1 (wt: Δ*spxA1*) co-culture is shown. (B) Cell perimeters of each condition were measured by Celltool and the data shown for the co-cultures indicate Δ*spxA1* cell perimeters. Data are from a single representative replicate where at least 1,000 cells or ten fields of view were quantified. The lines represent the means.

## SUPPLEMENTAL TABLES

Table S1. Proteins decreased in abundance in Δ*spxA1* compared to wt

Table S2. Proteins increased in abundance in Δ*spxA1* compared to wt

Table S3. Complete list of proteins detected by whole cell proteomics

Table S4. Strains used in this study

